# Growth, development, and life history of a mass-reared edible insect, *Gryllodes sigillatus* (Orthoptera: Gryllidae)

**DOI:** 10.1101/2024.10.08.617250

**Authors:** Jacinta D. Kong, Marshall W. Ritchie, Émile Vadboncoeur, Heath A. MacMillan, Susan M. Bertram

## Abstract

Insects provide a potential source of sustainable, alternative protein that can help meet the protein demands of a growing population. Efficient farming of insects to meet this demand depends on an understanding of insect life history. Yet, detailed information and expertise about a single species are not always available for practitioners to make informed decisions about rearing practices or identify arising issues. The cricket (*Gryllodes sigillatus*) is commonly farmed for human consumption or animal feed, but few studies have characterized the life history of this species throughout ontogeny. Here, we describe the growth and development of *G. sigillatus* from hatch to adulthood and quantify reproductive traits relevant to mass-rearing and colony management, including egg development. This information provides foundation to start and manage a cricket colony and to conduct research on growth and performance. We highlight ways that a fundamental understanding of cricket biology can be informative for optimizing cricket growth, reducing variability in yield and informing future precision farming practices.

## Introduction

Edible insects are a promising solution to the problem of feeding an increasing global population (Sorjonen et al. 2019, van Huis 2020). Insects have been economically and culturally important as food to many communities around the world (Raheem et al. 2019, Siddiqui et al. 2024). In recent years, commercial enterprises of intensive insect farming have gained attention as an alternative to conventional agriculture that aligns with the UN’s Sustainable Development Goals while meeting the demand for protein (UN 2015, Liceaga et al. 2022). Insect farming uses less space and water than conventional agriculture, is estimated to produce fewer greenhouse gasses, and can be integrated within sustainable practices, such as a circular economy (Chavez 2021, van Huis et al. 2021). To benefit from these advantages and to meet the world’s protein demand, the edible insect farming industry must continue to invest in generating actionable knowledge. Research and development play a crucial role in ensuring the edible insect industry develops effectively and sustainably by generating data that can inform, improve and optimize farming practices (Van Peer et al. 2024). Acquiring and implementing these data requires collaborative partnerships between academics, industry, and government agencies (Tomberlin et al. 2022, Robinson et al. 2024). Through these partnerships, research outcomes can be translated into commercial practice and can be used to quantitatively evaluate, refine, and adapt practices (Morales-Ramos et al. 2024).

Data-driven insect mass-rearing depends on fundamental biological information that draws expertise from agriculture and animal science, entomology, ecology, and evolution. Insects used in the edible insect industry are typically small, rapidly reproducing with short generation times and dense populations, and are amenable to manipulations of factors like temperature, diet, and humidity (Davidowitz 2021, Hawkey et al. 2021). Thus, the yield and productivity of commercial insect farms are highly dependent on species-specific biological traits that can be targets for research and development. Individual-level traits, such as instar progression, body size and mass throughout ontogeny, reproductive output and egg development, drive population dynamics that will determine the success of a colony and the resulting product yield. Variable yield among harvests is a problem that can emerge even when rearing conditions are unchanged (Moore and Martin 2019, Mutamiswa et al. 2023). Variability in yield generates uncertainty in the supply of insect products, which can slow the rate of industrial development needed for a profitable and sustainable industry (Larouche et al. 2023). One solution to this problem is to use a species-centric data-driven approach that allows for fine-scale optimization of rearing practice and for troubleshooting production issues. Examples of optimization studies in mass-reared insects include the effects of temperature for rearing (Chia et al. 2018, Mamai et al. 2018, Chen et al. 2022), diet preferences (Morales-Ramos et al. 2020), and reproduction and husbandry (Hoc et al. 2019). Failure to incorporate biological knowledge or make information publicly available can hinder efforts to troubleshoot production industry-wide (Larouche et al. 2023). However, the advantages of a data-driven farming approach also underpin the challenges of this approach; the requirements or costs for the data necessary to troubleshoot where and when in production variability in yield is generated can be prohibitively high (Davidowitz 2021). Therefore, publicly available fundamental science can play a key role in the continuing development of the insect farming industry and ensure consistent supply of products (Tomberlin et al. 2022).

Several true crickets (Orthoptera: Gryllidae) are farmed for consumption globally (Magara et al. 2021). In North America, the cricket *Gryllodes sigillatus* and the house cricket *Acheta domesticus* are the two main commercially farmed species (Larouche et al. 2023). *Acheta domesticus* has been widely used as a model species in entomology and more recently in an edible insect farming context because of its widespread use, leading towards calls for standardized data collection (Morales-Ramos et al. 2024, Van Peer et al. 2024). In contrast, *G. sigillatus* has received less attention particularly in a farming context despite its popularity as a farmed species and its long history of use as a model cricket species. There is a wealth of knowledge about the life history traits of adult *G. sigillatus* including digestive morphology (Biagio et al. 2009, Ritchie et al. 2024), immunity (Gershman et al. 2010), diet (Rapkin et al. 2018), lifespan (Archer et al. 2012), and reproductive effort to name a few (Bateman et al. 2001, Ketola and Kotiaho 2010, Okada et al. 2011). However, fewer studies have described the growth and development of *G. sigillatus* from hatch to adulthood and this information is directly relevant to farming (Muzzatti et al. 2022). Here, we aimed to fill this knowledge gap by describing the instar progression, growth, development, and egg production of a farmed strain of *G. sigillatus*. This information forms a fundamental basis for designing experiments for the optimization of rearing protocols for this species.

## Methods

### Colony care and husbandry

We maintained a colony of *Gryllodes sigillatus* at Carleton University, Ontario, Canada to ensure a consistent supply of individuals for several experiments. Separate cohorts of eggs from this colony were used to quantify life history characteristics in this study. These individuals were originally sourced as eggs from Entomo Farms, Ontario, Canada that have been mass-rearing a population for food and feed for over a decade. The colony was maintained at 30°C (30.24 ± 1.43°C, mean ± standard deviation), 30% relative humidity (R.H.) (30.88 ± 11.01%), and a 14:10 h Light:Dark photoperiod. Room temperature and R.H. were monitored using data loggers (IBS-TH2, Inkbird, Guangdong, China). All crickets were provided food (cricket feed mix of soy, corn, fishmeal and micronutrients, 1:0.94 Protein: Carbohydrate, Campbell Feed Mill, Ontario, Canada), water as wet paper towels in a vial, and egg carton as shelter. The colony consisted of ten 5.2 L plastic containers (22.9 × 15.2 × 15.2 cm, L × W × H) containing approximately 200 crickets in each container until instars 6-7, then the number of crickets was reduced to 20 per container to minimize potential effects of high rearing density. As part of the continued colony maintenance, adult crickets were allowed to oviposit into moist coco peat for 48 h. The peat was collected and incubated under the same conditions as the parents until the eggs within hatched. Egg development time was taken as the time between when the peat was added to the adult colony and the day the first hatched nymph was observed. The hatched nymphs were either used to start a cohort to be used in experiments or another generation of the colony. Each container of the next generation of the colony was seeded from two parental containers to allow genetic mixing.

### Nymph growth, development and morphometrics

To capture instar progression, a cohort of ∼500 crickets from the colony were housed in a plastic bin (68 L, 61 x 41 x 32 cm) under colony conditions. Crickets were randomly sampled every 2-3 days for seven weeks (355 individuals in total). Individual crickets were anesthetized with CO_2_ for 15 s, weighed to obtain wet mass, and then photographed under a dissecting microscope (Stemi 508 with an axiocam 105 color, Zeiss, Oberkochen, Germany). Dorsal and ventral photographs were obtained for each individual. Sex if possible was determined through visual inspection of the photo, and the body parts were measured in Fiji (ImageJ) v2.3.051 (Schindelin et al. 2012). Body parts measured were maximum head width (distance between the two outside edges of the eyes), maximum thorax width, maximum thorax length, maximum abdomen width, maximum abdomen length and maximum femur length (**Fig. 1**). A subset of these crickets were then frozen at -20°C (n = 265), then desiccated in an oven at 60°C for at least 72 h (Precision Scientific, IL, U.S.A.), and then weighed using a microbalance (ME5, Sartorius, Göttingen, Germany) to determine dry mass and calculate water content.

**Fig. 1.**
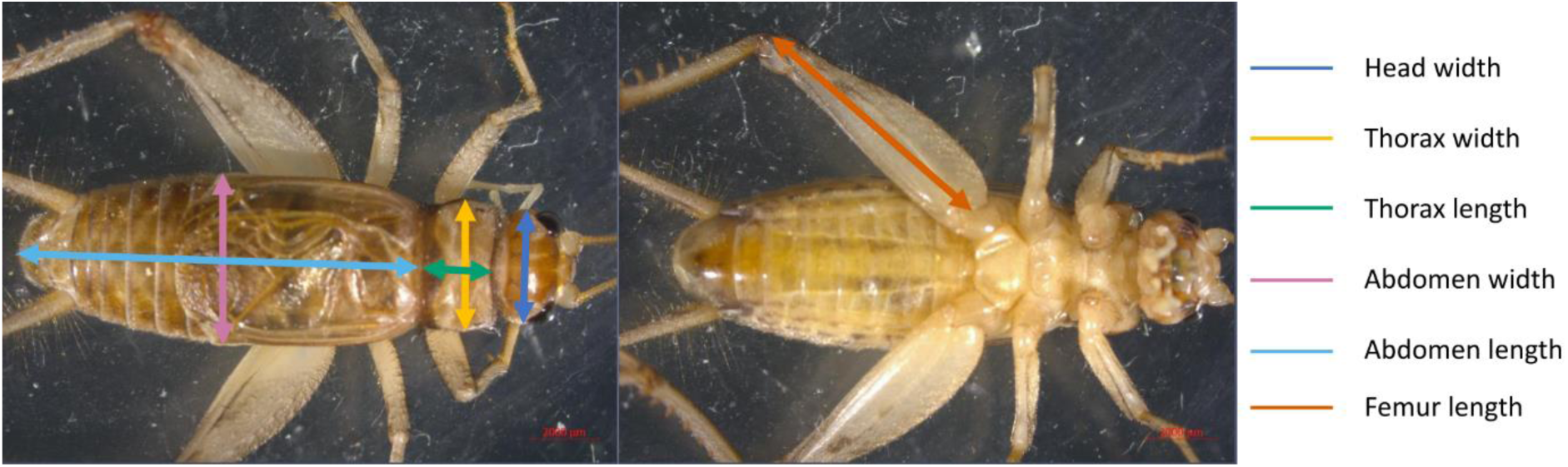
Morphometric landmarks for *Gryllodes sigillatus* from dorsal (left) and ventral (right) photographs. Adult eighth instar male shown. Images were analyzed in ImageJ.

### Fecundity

A cohort of ∼100 crickets originating as eggs from the colony were kept together from hatch until sex could be easily determined (6^th^ instar), then assigned separate containers for males and females. When crickets had reached adulthood (8^th^ instar, 39 days since first nymphs hatched), male and female pairs were set up and kept under colony conditions (31.15 ± 1.73°C and 33.56 ± 4.95% R.H., mean ± standard deviation). Temperature and relative humidity was monitored using a data logger on the same shelf as the crickets for the duration of the experiment (IBS-TH2, Inkbird, Guangdong, China). Crickets were weighed (AB135-S, Mettler Toledo, ON, Canada) to obtain adult mass then randomly assigned mating pairs. Mating pairs were housed in a food container (710 mL, 190 x 127 x 70 mm) with a piece of egg carton for shelter and fresh feed in an ink cap (17mm diameter). Fresh feed was replaced every week. A small container filled with marbles and water (96 mL, 70 mm diameter x 30 mm) provided water and an oviposition substrate. Water was chosen as an oviposition substrate as crickets will readily oviposit in water and it is easier to remove eggs for counting than sand or peat, but crickets may drown. Drowning accounted for most of the early mortality in this study. Eggs were decanted from the marbles and water every two days, at which time the water was refilled. Photographs of egg batches were used to count large batches. Males were removed if the paired female died. Females were retained if the paired male died. Eggs were collected until all surviving crickets had oviposited eggs for 30 days (38 days in total since the start of pairing). The delay in first oviposition was calculated as the difference between when the first batch of eggs was oviposited and when crickets were allocated to mating pairs. Fecundity was calculated as the mean number of eggs oviposited every 48 h over the 30-day period since first oviposition for each pair. Two pairs of the original 25 pairs were excluded from analysis because one pair died two days into the experiment without ovipositing and one pair died after ovipositing a single batch of egg on day six, giving 23 pairs for analysis.

### Statistical analysis

Data were analyzed in R v4.3 (R Core Team 2023). To assess autocorrelation amongst closely related traits, we calculated Pearson correlation coefficients for pairwise morphometric traits, pooling sex (n = 355). To assess overall relationships between morphological traits and instar, we fitted linear regressions to Log_10_ transformed morphological traits, pooling sex. Morphological traits were Log_10_ transformed to meet the assumption of linearity. To assess for sex-specific differences in morphological traits, a separate linear regression was fitted to the subset of data for instars 5-8 when sex could be identified, and morphological traits were not Log_10_ transformed.

To account for heteroskedasticity in wet mass as crickets grow, we fitted a weighted logistic growth curve to wet mass over time with a power variance function for weighting wet mass using *nlme* v3.1 (Pinheiro and Bates 2009). An initial likelihood ratio test showed the weighted logistic regression provided a better fit to the data than the unweighted regression (Likelihood ratio = 982.7, P < 0.001), which showed heteroskedasticity in standardized residuals. Relationships between wet mass and instar was Log_10_ transformed to meet the assumption of linearity. Mass and instar were not included as predictor variables in the same model to avoid autocorrelation (r = 0.95). We used linear regression to assess the relationship between untransformed wet and dry mass.

Fecundity was analyzed with a mixed effects model using *nlme* with time standardized to first oviposition as a fixed continuous variable and pair as a random variable (n = 23). Time was standardized to account for the variation in when oviposition starts among pairs. The effect of time on fecundity was analyzed using a Wald test and the random effect of pair on fecundity was tested using a likelihood ratio test (Analysis of Deviance) against the reduced fixed effects model.

## Results

### Egg development

The first nymphs emerged at similar times among independent containers (**Table 1**). Emergence during the first 24 h was minimal and peaked over the next 48 h. Emergence reduced three days after the first nymphs emerged and we observed this trend across independent containers even though emergence over time was not formally quantified.

**Table 1.** Life history traits of mass-reared crickets, *Gryllodes sigillatus*. Crickets were reared at 30°C.

| Trait | Mean value ± standard deviation (sample size) |
| --- | --- |
| Egg development time (days) | 10.6 ± 0.5 (20) |
| Mean time to first oviposition (days) | 5.66 ± 1.67 (23) |
| Fecundity (Mean number of eggs per 48 h per female within 30 days of oviposition) | 56.74 ± 31.77 (23) |
| Dry mass at hatch (24h old nymphs, mg) | 0.0313 ± 0.016 (9) |

### Morphometrics

The life cycle of *G. sigillatus* consisted of eight instars that are identifiable by morphometric characteristics (**Fig. 1**). Crickets molted every 3-4 days between instars 1 to 5 in the first two weeks. These instars are identifiable in photographs by their relative size and shape but lacked distinguishing characteristics easily visible to the naked eye (**Fig. 2**).

**Fig. 2.**
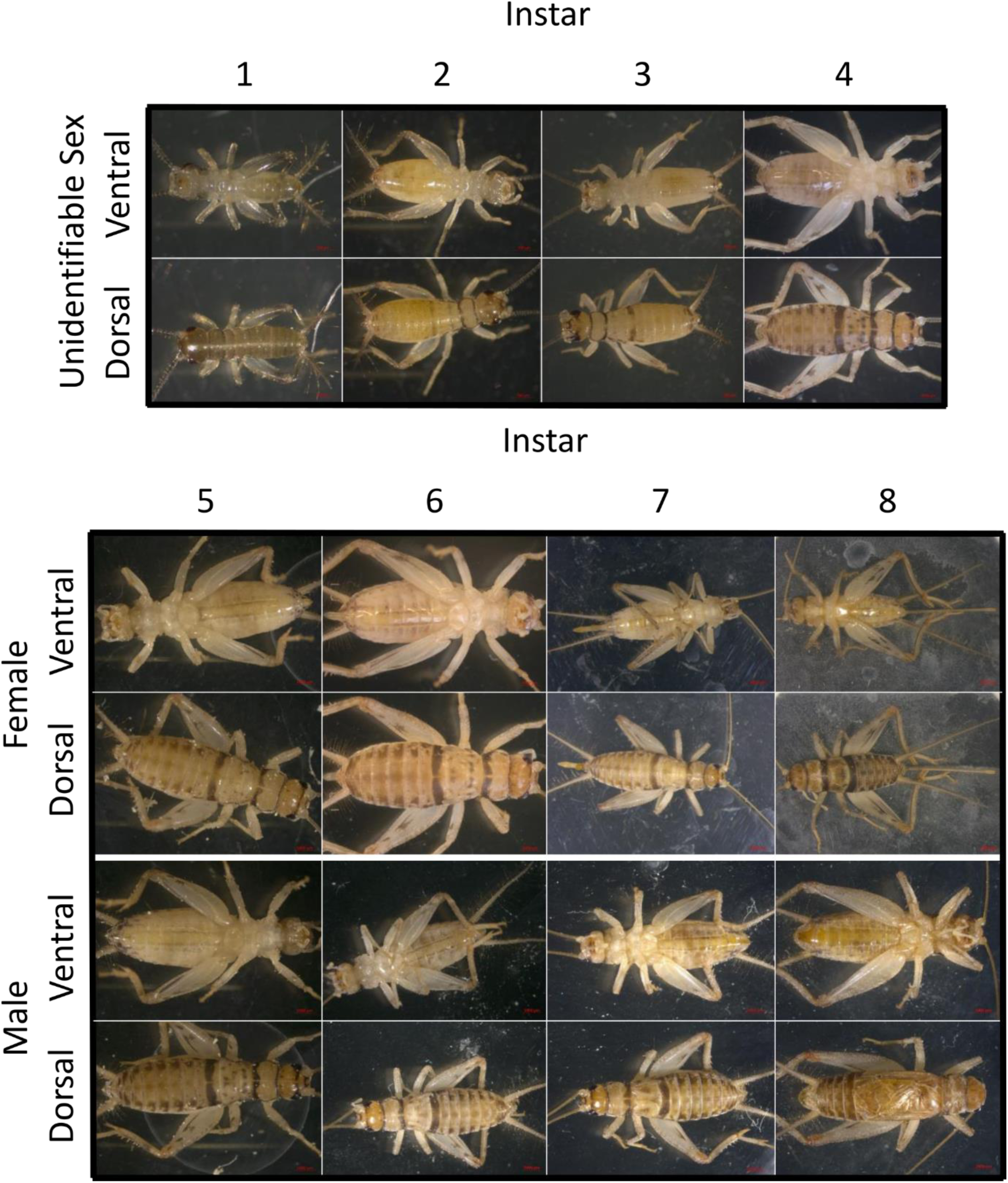
Ventral and dorsal photos of representative *Gryllodes sigillatus* crickets for each instar (columns) and sex (Unidentifiable, Female and Male).

Sex was first identifiable at instar 5 when ovipositors first appeared in females, but because the ovipositor does not extend beyond the abdomen and it was not easily visible without magnification (**Fig. 2**). Ovipositor length of instar 6 and 7 females varied among individuals of the same instar but were visually distinctive and distinct from a fully formed ovipositor of instar 8. Wing buds on females first appeared at instar 7, while they appeared on males at instar 6. The size and presence of wing buds were variable, particularly in females, but were distinctive for males of instar 6 and 7 and were absent in instar 5 males. Crickets reached reproductive maturity at instar 8 (adult stage), characterized by fully formed ovipositors in females and wings in males. Males started signaling acoustically for mates at instar 8. Instars 6 to 8 were easily identifiable by these sex-specific morphological characteristics but were less easily distinguished by size and shape alone as data clusters of instars overlapped (**Fig. 2**, **Fig. 3**).

**Fig. 3.**
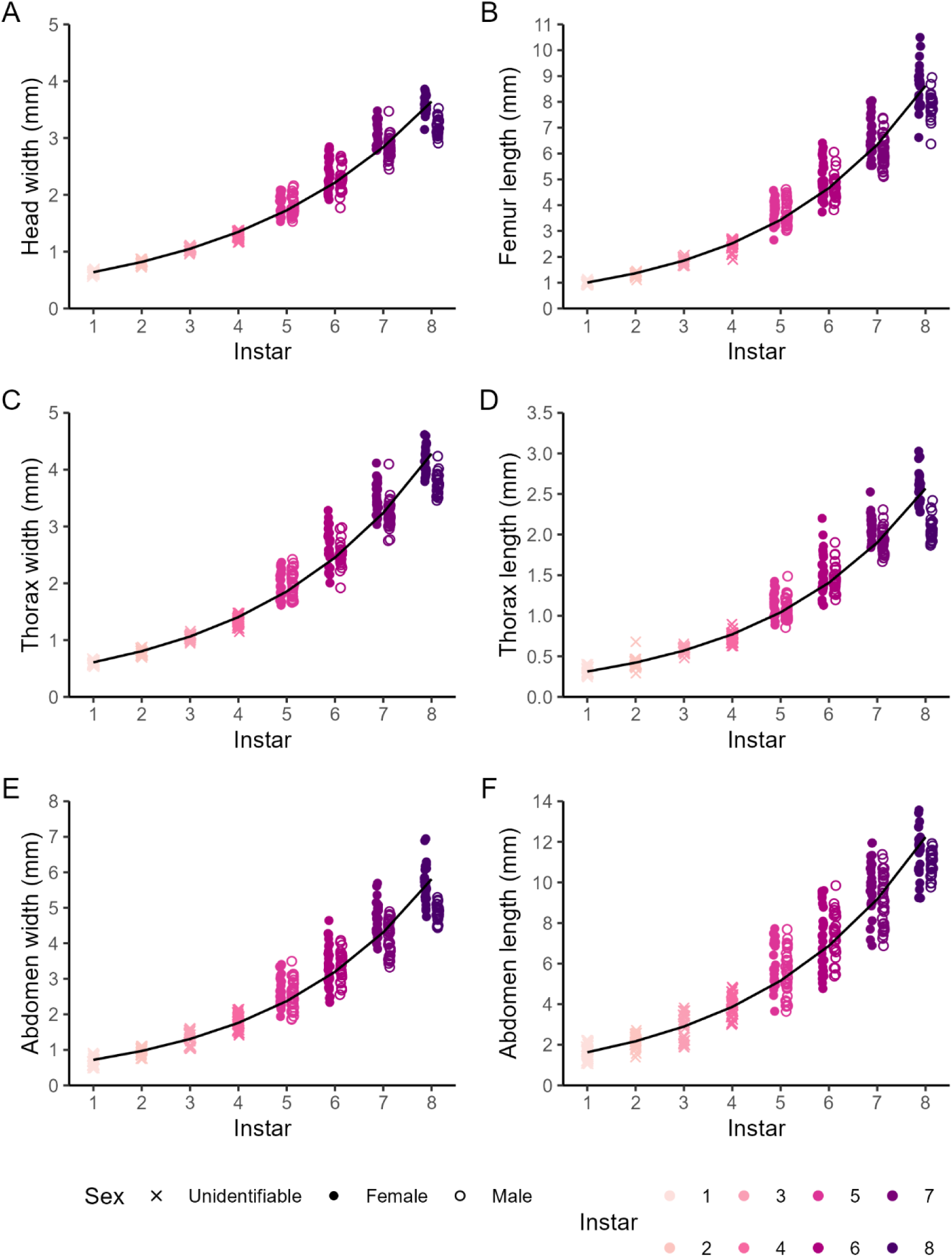
A) Head width, B) Femur length, C) Thorax width, D) Thorax length, E) Abdomen width, and F) Abdomen length for each instar of *Gryllodes sigillatus* (colors, n = 355). Crosses denote juvenile crickets of unidentifiable sex (instars 1 – 4), closed circles denote female crickets (instars 5 – 8), and open circles denote male crickets (instars 5 – 8). Solid black line indicates the fitted linear regression to Log_10_ transformed traits, pooling sex (**Table 2**).

**Table 2.** Regression equations and Coefficient of Determination, R^2^ (%), for relationships shown in. **Figs. 3 and 4**.

| Figure panel | Trait | Equation | $R^2$ (%) |
| --- | --- | --- | --- |
| **Fig. 3A** | Head width (mm) by instar | $\text{Log}_{10}(\text{Head width}) = 0.11 \times \text{Instar} - 0.30$ | 97.59 |
| **Fig. 3B** | Femur length (mm) by instar | $\text{Log}_{10}(\text{Femur length}) = 0.13 \times \text{Instar} - 0.13$ | 97.52 |
| **Fig. 3C** | Thorax width (mm) by instar | $\text{Log}_{10}(\text{Thorax width}) = 0.12 \times \text{Instar} - 0.34$ | 97.57 |
| **Fig. 3D** | Thorax length (mm) by instar | $\text{Log}_{10}(\text{Thorax length}) = 0.13 \times \text{Instar} - 0.64$ | 96.41 |
| **Fig. 3E** | Abdomen width (mm) by instar | $\text{Log}_{10}(\text{Abdomen width}) = 0.13 \times \text{Instar} - 0.27$ | 94.90 |
| **Fig. 3F** | Abdomen length (mm) by instar | $\text{Log}_{10}(\text{Abdomen length}) = 0.13 \times \text{Instar} + 0.09$ | 92.25 |
| **Fig. 4A** | Wet mass (mg) over time | $\text{Wet mass, mg (time)} = \frac{237.61}{1 + e^{\left(\frac{25.38 - \text{time}}{4.35}\right)}}$ | - |
| **Fig. 4B** | Wet mass (mg) by instar | $\text{Log}_{10}(\text{Wet mass}) = 0.35 \times \text{Instar} - 0.30$ | 95.57 |
| **Fig. 4C** | Wet mass (mg) by dry mass (mg) | $\text{Wet mass} = 3.17 \times \text{Dry mass} + 8.72$ | 97.10 |

Transformed morphometric traits followed tight linear correlations with instar where data clustered at each instar (**Fig. 3**). Changes in morphometric traits with increasing wet mass was best described by a power function (**Supp. Fig. S1**, **Supp. Table S1**) and all morphometric traits are highly correlated with each other (*r* = 0.95 – 1, **Supp. Fig. S2**). Head width was the most reliable single predictor of instar, as it increased stepwise throughout ontogeny and was relatively invariant within an instar (**Table 2**). Head width also positively correlated with mass and had the highest coefficient of determination of all the morphological traits we measured. As females were larger overall than males, females had larger head widths than males and larger increases over time (Instar × Sex: t_227_ = −4.19, P < 0.001).

### Growth curve and dry mass

Increases in wet mass followed a sigmoidal pattern throughout ontogeny (**Table 2**, **Fig. 4A**) and increased exponentially with instar progression (**Table 2**, **Fig. 4B**). Dry mass had a tight relationship with wet mass (**Table 2**, **Fig. 4C**). We observed sexual size dimorphism at instars 7 and 8, but not earlier. Females were heavier and had larger increases in mass over time than males (**Fig. S3**, Instar × Sex: t_227_ = −6.78, P < 0.001). Crickets had a water content of 75.7% on average across all instars, and adult females had a lower water content than adult males (**Table 3**).

**Fig. 4.**
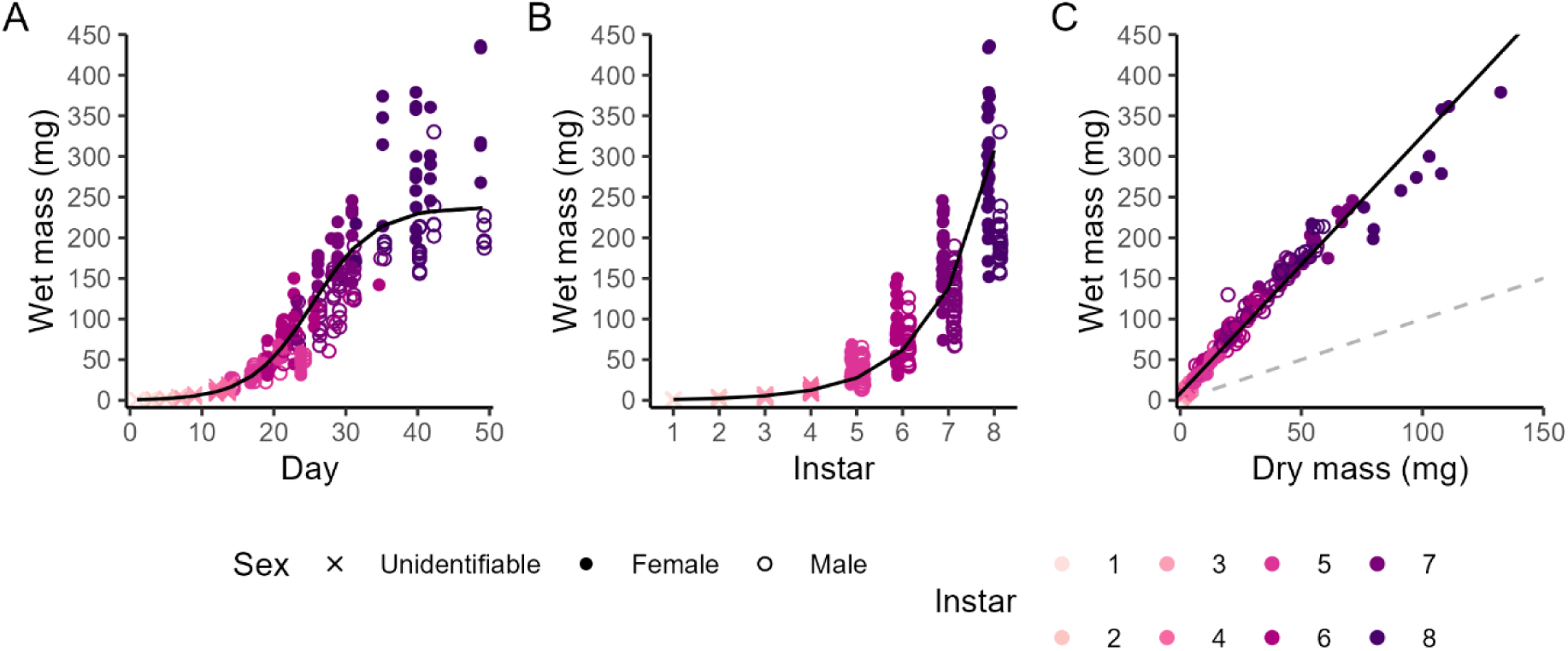
Cricket (*Gryllodes sigillatus*) mass for A) wet mass over time (n = 355), B) wet mass by instar (colors), and C) wet mass against dry mass (mg, n = 265) Crosses denote juvenile crickets of unidentifiable sex, closed circles denote female crickets, and open circles denote male crickets. Dashed grey line indicates the identity line and the solid black line indicates the fitted regressions (**Table 2**).

**Table 3.**
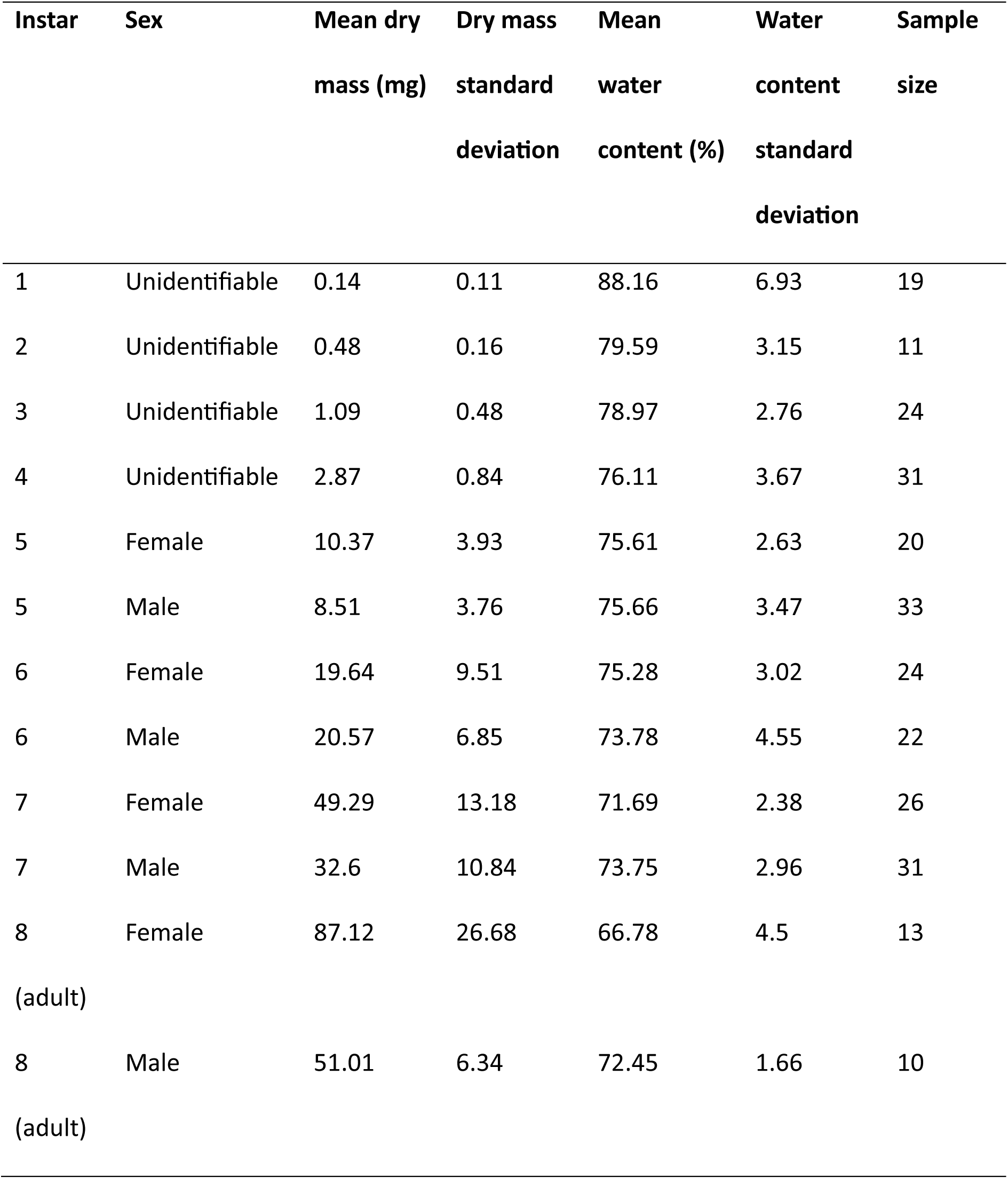
Dry mass (mg) and water content (%) of crickets (*Gryllodes sigillatus*) throughout ontogeny.

### Fecundity

We observed some crickets mated immediately after pairing but we did not quantify mating behavior. The first eggs were oviposited between 2 and 10 days after pairing (median 6 days, **Table 2**). Fecundity of a pair (mean number of eggs oviposited in each 48h period within 30 days of the start of oviposition, n = 23) ranged from 0 to 101.67 eggs (**Table 1**, **Fig. 5**). One pair did not oviposit any eggs for 18 days until the female died. The maximum number of eggs oviposited each 48h period ranged from 0 to 279 eggs across the mated pairs. Not all pairs oviposited eggs every two days (**Fig. 5C**; cumulative increases in the number of eggs oviposited was not linear). Overall mean fecundity did not change over time (t_1, 250_ = –1.95, P = 0.052) and trends in fecundity varied among pairs (χ^2^ = 13.62, P < 0.001). Females continued to oviposit eggs after their male partner had died. Five of the 15 pairs remaining at the end of the experiment were a single female. Damaged ovipositors were observed in some females over time and most females appeared to stop ovipositing after this damage, but one female was observed to oviposit with a damaged ovipositor.

**Fig. 5.**
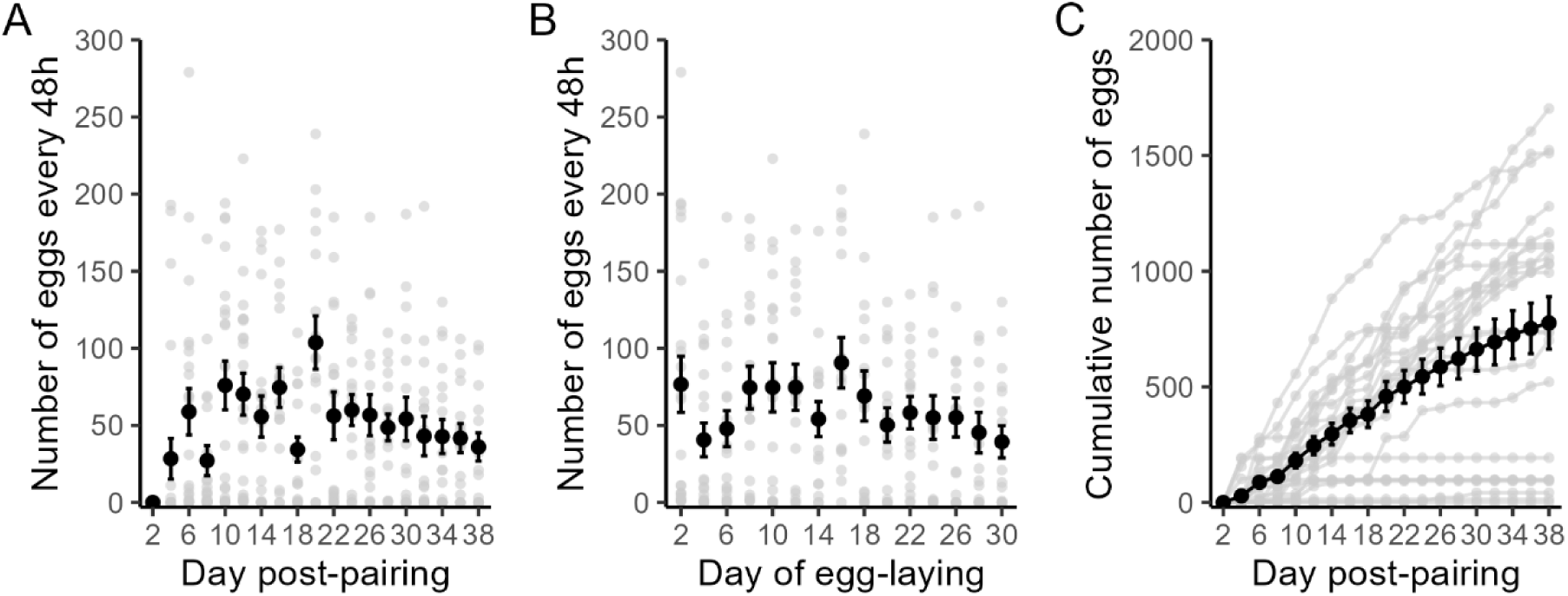
Fecundity (number of eggs every 48 hours) of male and female pairs over time from A) when crickets (*Gryllodes sigillatus*) were paired until the end of the experiment, and B) from first oviposition until 30 days had passed. C) Cumulative number of eggs for each pair. Grey points indicate each batch of eggs from each pair (n = 23) and black points and bars represent the mean and standard error across pairs.

## Discussion

An in-depth understanding of the biology and life history of a farmed species provides a foundation for understanding the drivers and determinants of variation. This information can be leveraged to optimize farming practices and troubleshoot variation in yields (Davidowitz 2021). However, the high-resolution information needed to make these inferences takes time, labor, and expertise to collect, and these data are not always publicly available (Tomberlin et al. 2022, Larouche et al. 2023). Here, we aimed to describe the life history of a commonly farmed cricket *Gryllodes sigillatus*. We characterized morphological progression through the instars, changes in wet and dry mass as well as water content throughout ontogeny, and fecundity. This information provides a basis for designing manipulative studies into the allometry, growth, development and reproduction of this species that can target performance traits important to mass-rearing of this species.

We found *G. sigillatus* instars are identifiable, especially late nymphal stages between when sexual characteristics first appear and the adult stage. Instars could be estimated from expert inspection and estimates validated from trait values. We observed that the condition of the individual adds uncertainty in estimating instar because morphological measurements vary within instars. Specifically, abdomen length was a less reliable indicator of instar or body size because the abdomen may greatly increase in length and width after eating (Deku et al. 2022). Thorax width may also extend after eating, but to a lesser extent than the abdomen length. However, evaluating morphological traits depends on intended use. For example, abdomen size may be relevant for traits associated with egg production or feeding (Fudlosid et al. 2022). Head width provided the best discrimination for instar, is used to discriminate instars in other insect species (Calvo and Molina 2008), and is highly correlated with other morphological traits (**Fig. S2**). However, we suggest using two or more morphometric traits in combination with wet mass to obtain the most accurate estimation of instar, especially for instars 1 – 4, as the overlap of data between close instars may lead to incorrect estimations, particularly for individuals that deviate from the mean.

Wet mass was highly correlated with morphological traits but was more variable than morphology for estimating instar. This pattern is driven by larger variation in wet mass within an instar (**Supp. Fig. S1**, individuals of the same instar and sex fall along the x-axis with minimal change in morphological trait values). The decoupling of wet mass and body size through instar progression has been reported across insects (Chown and Gaston 2010). One reason for this decoupling is the change in body composition from the development of sexual characteristics at later instars. For example, adult females had a lower water content than adult males that likely reflects the development of fatty tissues such as eggs and ovaries (**Table 3**). Changes in internal anatomy, such as the presence of extensive air sacs, which are necessary to support large body sizes do not increase body mass proportionally (Harrison et al. 2023). This overall phenomenon is relevant to insects farmed for food and feed because desirable nutritional qualities for a final product may not tightly correlate with body mass. That is, increases in body size may not increase the nutritional content of a cricket because its larger body size is attributed to a more extensive tracheal system required to maintain a larger body size. Thus, selection on body size and not body mass may not be beneficial for a farm. Crickets that are heavier may contain more tissue that contributes to the final nutritional content of the product. Our body water content estimates were higher than other Orthopterans but water content varies greatly among insect species (Hadley 1994, Woodman 2012).

Our study used instar as a proxy for developmental stage rather than time as other studies have done. Few studies have reported growth and development of *G. sigillatus* over ontogeny and these studies have used week as a proxy for age and instar (e.g. McFarlane 1964, Fudlosid et al. 2022, Muzzatti et al. 2022). Other studies are limited to later nymphal stages or the adult stage (e.g. Rapkin et al. 2018). Our data can be used to infer instar from studies reporting age in time or from wet mass or body size. A fine-scale understanding of growth allows for certain time periods that may be of interest for a farming context to be targeted (Morales-Ramos et al. 2015). For example, the fastest growth estimated by the logistic model occurs between instars 6 and 7 (approximately day 25, **Table 2**) which is when sexual morphological traits are developing (**Fig. 4A**). This transitional period may be of interest to understand patterns of maturation or responses to stressors over ontogeny that manifest at adulthood (Nowosielski and Patton 1965, Klockmann et al. 2017b). As the growth pattern of this species is sigmoidal, which is typical of insects, a sampling interval of weeks may not be able to identify or capture this critical period of growth. The timing of the inflection point in the growth curve may also change depending on the individual and the rearing conditions. Inter-individual variation in maturation in *G. sigillatus* can be high during this period of maximum growth and as sexual dimorphism emerges, even under controlled conditions. Being able to identify instar is important for ensuring sampling consistency and quantifying this inter-individual variation (much more overlap in data points in **Fig. 4A** than in earlier timepoints). For example, studies that aim to target the adult stage, such as for reproduction, will benefit from being able to identify when a cricket matures to control for the age of the cricket. Understanding the physiological basis for growth and maturation is a fundamental biological problem that encompasses allometry, drivers of optimal growth, and to what extent growth and maturation are fixed for an individual (Tennessen and Thummel 2011, Mirth et al. 2016, Meister et al. 2017, Buchanan et al. 2022). These fundamental problems can be directly translated to commercially farmed insects. From a farming perspective, variation in maturation affects the logistics around when eggs can be collected from a breeding population and for how long, as well as the timing of a harvest.

Our understanding of *G. sigillatus* phenotypes and life history complement several challenges of mass-rearing crickets. First, genomics represents an untapped potential for farmed insects and our ability to link genes with fitness related phenotypes that are beneficial for farming, or to target specific genes, depends on characterizing life history phenotypes (Nakamura et al. 2022). *Gryllodes sigillatus* has a tropical worldwide distribution including in urban areas and human habitats, which likely permits range extensions far into temperate areas through the movement of individuals for the pet food industry (Masaki 1978, Toms 1993, Bateman et al. 2005). This distribution suggests *G. sigillatus* is adaptable to a range of habitats within their environmental tolerance. However, most studies on *G. sigillatus,* including the present study, use laboratory or farm populations that have been removed from their wild type ancestors for several generations and years. It is likely there is significant variation in traits among populations or strains that have evolved through the separation of farmed populations over time. For examples, our mean fecundity in a 48h period was lower than previously reported for outbred lines of *G. sigillatus* for the same sampling period, we did not find a decrease in fecundity over time (as a proxy for parental age) as previously reported for a similar time frame (30 days) (Archer et al. 2012), and longevity and resting metabolic rate differed between two laboratory populations of *G. sigillatus* sourced from USA and Australia (Okada et al. 2011). Studies on inbred lines of *G. sigillatus* demonstrated that sustained inbreeding has profound negative effects on life history traits and fitness that would affect colony maintenance, such as decreased female fecundity, decreased male calling durations, and decreased immune responses to an immunity challenge (Gershman et al. 2010, Sakaluk et al. 2019). These pedigree studies also suggest that *G. sigillatus* would be responsive to directed selection and experimental evolution. Thus, there is potential to explore the development of strains optimized for mass-rearing which may also minimize variability in yields, while considering that selection on body size may not correspond with body mass or desired product qualities (Larouche et al. 2023).

Second, disease is an important issue for mass-rearing insects from an animal welfare and food safety perspective (Fernandez-Cassi et al. 2020, Vogel et al. 2022). Viruses, parasites, and microbes that negatively affect the performance of mass-reared insects may contribute towards variability in farming yields (Bertola and Mutinelli 2021, Jordan and Tomberlin 2021, Herren et al. 2023). To monitor and decide when to intervene in a population requires an understanding of what a healthy population looks like, and what indicators of stress precede a potential colony collapse. In this context, life history data as well as behavioral data are valuable for training artificial intelligence models to monitor insect welfare in mass-reared contexts through changes in behavioral patterns or visual signals (Hansen et al. 2022, Wenning et al. 2022), and can complement genomic approaches for monitoring disease (Lim et al. 2024).

Third, external variables such as temperature, desiccation, density and food availability during the egg or early nymphal stages even as a short-term acute stress can have profound and lasting consequences on insect performance (Klockmann et al. 2017a, Mahavidanage et al. 2023). For example, a simulated heat wave during the juvenile stage can reduce reproductive fitness in an insect (Stahlschmidt et al. 2023). These consequences of early life stress are well documented for holometabolous insects but are not as well documented for hemimetabolous insects, like crickets (Urquhart-Cronish and Sokolowski 2014, Zhang et al. 2015, Klockmann and Fischer 2017). Egg and early nymph stages are often handled differently to adults in an insect farm, sometimes in a separate facility under different environmental conditions. There are opportunities to investigate whether these differences in handling can explain variability in yields and design best practices or decide when to intervene to ensure consistency in yields (Morales-Ramos et al. 2024).

Life history provides a framework in which to understand variation in performance that in turn underpins yield. Knowledge of instars is useful for tracking progression of development, identifying important stages of development, and assessing inter-individual variation in a population (Bien et al. 2023). More generally, high resolution morphological data and validated instar estimates are essential for training artificial intelligence models that aim to record growth and development of individual crickets, as well as population sizes in high density precision farming, as have been developed for other livestock (Tedeschi et al. 2021, Bao and Xie 2022). To fully leverage these emerging technologies, the accuracy and precision of the machine learning model needs to be assessed under less controlled conditions such as conditions expected on a farm, and on datasets ranging in quality (Wenning et al. 2022). Moving beyond monitoring, our understanding of insect growth and development directly informs models of animal growth that can be used to predict growth and the timing of optimal harvests (Mauritsson and Jonsson 2023). For example, life history traits, feeding rates, and metabolic rate can feed into a Dynamic Energy Budget model to predict growth and development of an insect. These models exist for holometabolous insects but have only recently been developed for a hemimetabolous insect (Llandres et al. 2015, Klagkou et al. 2024). Overall, an integrative and collaborative approach between stakeholders will facilitate research breakthroughs in insect farming for food and feed (Robinson et al. 2024).

## Acknowledgements

We thank Matthew Muzzatti and Nirush Arulendran for assistance with animal care and data collection, and to Entomo Farms for providing crickets and materials for this study. This study was supported by an NSERC-Mitacs alliance grant (ALLRP 568647 – 21) awarded to S. M. B. and H. A. M. in partnership with Entomo Farms and Aspire Food Group, Canada. It was further supported by an NSERC Discovery Grants (RGPIN-2018-05322 and RGPIN-2017-06263) and infrastructure supports from the Canadian Foundation for Innovation to H.A.M. and S.M.B., respectively.

## Author contributions

JDK: Conceptualization; Data curation; Formal analysis; Investigation; Methodology; Project administration; Visualization; Writing—original draft; Writing—review & editing. MWR: Investigation; Methodology; Writing—original draft; Writing—review & editing. ÉV: Investigation; Methodology; Writing— review & editing. HAM: Funding acquisition; Supervision; Writing—review & editing. SMB: Funding acquisition; Supervision; Writing— review & editing.

## References

Archer CR, Zajitschek F, Sakaluk SK et al. 2012. Sexual selection affects the evolution of lifespan and ageing in the decorated cricket *Gryllodes sigillatus*. Evolution 66(10):3088–3100. 10.1111/j.1558-5646.2012.01673.x.

Bao J, Xie Q. 2022. Artificial intelligence in animal farming: A systematic literature review. Journal of Cleaner Production 331(129956). 10.1016/j.jclepro.2021.129956.

Bateman PW, Gilson LN, Ferguson JWH. 2001. Investment in mate guarding may compensate for constraints on ejaculate production in the cricket *Gryllodes sigillatus*. Ethology 107(12):1087–1098. 10.1046/j.1439-0310.2001.00756.x.

Bateman PW, Verburgt L, Ferguson JWH. 2005. Exposure to male song increases rate of egg development in the cricket *Gryllodes sigillatus*. Afr Zool 40(2):323–326.

Bertola M, Mutinelli F. 2021. A systematic review on viruses in mass-reared edible insect species. Viruses-Basel 13(11). 10.3390/v13112280.

Biagio FP, Tamaki FK, Terra WR et al. 2009. Digestive morphophysiology of *Gryllodes sigillatus* (Orthoptera: Gryllidae). J Insect Physiol 55(12):1125–1133. 10.1016/j.jinsphys.2009.08.015.

Bien T, Alexander BH, White E et al. 2023. Sizing up spotted lanternfly nymphs for instar determination and growth allometry. PLOS ONE 18(2):e0265707. 10.1371/journal.pone.0265707.

Buchanan KL, Meillère A, Jessop TS. 2022. Early life nutrition and the programming of the phenotype’. In: Costantini D, Marasco V *eds.* *Cham*: *Springer International Publishing*. 161*-*214.

Calvo D, Molina JM. 2008. Head capsule width and instar determination for larvae of *Streblote panda* (Lepidoptera: Lasiocampidae). Ann Entomol Soc Am 101(5):881–886. 10.1603/0013-8746(2008)101[881:HCWAID]2.0.CO;2.

Chavez M. 2021. The sustainability of industrial insect mass rearing for food and feed production: zero waste goals through by-product utilization. Curr Opin Insect Sci 48:44–49. 10.1016/j.cois.2021.09.003.

Chen L, Sørensen JG, Enkegaard A. 2022. Acclimation for optimisation: effects of temperature on development, reproduction and size of *Trichogramma achaeae*. Biocontrol Sci Technol 32(1):60–73. 10.1080/09583157.2021.1963679.

Chia SY, Tanga CM, Khamis FM et al. 2018. Threshold temperatures and thermal requirements of black soldier fly *Hermetia illucens*: Implications for mass production. PLOS ONE 13(11):e0206097. 10.1371/journal.pone.0206097.

Chown SL, Gaston KJ. 2010. Body size variation in insects: a macroecological perspective. Biological Reviews 85(1):139–169. 10.1111/j.1469-185X.2009.00097.x.

Davidowitz G. 2021. Habitat-centric versus species-centric approaches to edible insects for food and feed. Curr Opin Insect Sci 48:37–43. 10.1016/j.cois.2021.09.006.

Deku G, Combey R, Doggett SL. 2022. Morphometrics of the tropical bed bug (Hemiptera: Cimicidae) from Cape Coast, Ghana. J Med Entomol 59(5):1534–1547. 10.1093/jme/tjac072.

Fernandez-Cassi X, Söderqvist K, Bakeeva A et al. 2020. Microbial communities and food safety aspects of crickets (*Acheta domesticus*) reared under controlled conditions. Journal of Insects as Food and Feed 6(4):429–440. 10.3920/JIFF2019.0048.

Fudlosid S, Ritchie MW, Muzzatti MJ et al. 2022. Ingestion of microplastic fibres, but not microplastic beads, impacts growth rates in the tropical house cricket *Gryllodes sigillatus*. Front. Physiol. 13(10.3389/fphys.2022.871149.

Gershman SN, Barnett CA, Pettinger AM et al. 2010. Inbred decorated crickets exhibit higher measures of macroparasitic immunity than outbred individuals. Heredity 105(3):282–289. 10.1038/hdy.2010.1.

Hadley NF. 1994. Water relations of terrestrial arthropods. *San Diego*: Academic Press, Inc.

Hansen MF, Oparaeke A, Gallagher R et al. 2022. Towards Machine Vision for Insect Welfare Monitoring and Behavioural Insights. Frontiers in Veterinary Science 9(10.3389/fvets.2022.835529.

Harrison JF, McKenzie EKG, Talal S et al. 2023. Air sacs are a key adaptive trait of the insect respiratory system. J Exp Biol 226(10):jeb245712. 10.1242/jeb.245712.

Hawkey KJ, Lopez-Viso C, Brameld JM et al. 2021. Insects: A potential source of protein and other nutrients for feed and food. Annual Review of Animal Biosciences, Vol 9, 2021 9 333–354. 10.1146/annurev-animal-021419-083930.

Herren P, Hesketh H, Meyling NV et al. 2023. Environment–host–parasite interactions in mass-reared insects. Trends Parasitol 39(7):588–602. 10.1016/j.pt.2023.04.007.

Hoc B, Noël G, Carpentier J et al. 2019. Optimization of black soldier fly (*Hermetia illucens*) artificial reproduction. PLOS ONE 14(4):e0216160. 10.1371/journal.pone.0216160.

Jordan HR, Tomberlin JK. 2021. Microbial influence on reproduction, conversion, and growth of mass produced insects. Curr Opin Insect Sci 48:57–63. 10.1016/j.cois.2021.10.001.

Ketola T, Kotiaho JS. 2010. Inbreeding, energy use and sexual signaling. Evol Ecol 24(4):761–772. 10.1007/s10682-009-9333-1.

Klagkou E, Gergs A, Baden CU et al. 2024. Dynamic Energy Budget approach for modeling growth and reproduction of Neotropical stink bugs. Ecol Modell 493(110740. 10.1016/j.ecolmodel.2024.110740.

Klockmann M, Fischer K. 2017. Effects of temperature and drought on early life stages in three species of butterflies: Mortality of early life stages as a key determinant of vulnerability to climate change? Ecol. Evol. 7(24):10871–10879. 10.1002/ece3.3588.

Klockmann M, Kleinschmidt F, Fischer K. 2017a. Carried over: Heat stress in the egg stage reduces subsequent performance in a butterfly. PLOS ONE 12(7):e0180968. 10.1371/journal.pone.0180968.

Klockmann M, Günter F, Fischer K. 2017b. Heat resistance throughout ontogeny: body size constrains thermal tolerance. Glob Change Biol 23(2):686–696. 10.1111/gcb.13407.

Larouche J, Campbell B, Hénault-Éthier L et al. 2023. The edible insect sector in Canada and the United States. Animal Frontiers 13(4):16–25. 10.1093/af/vfad047.

Liceaga AM, Aguilar-Toalá JE, Vallejo-Cordoba B et al. 2022. Insects as an alternative protein source. Annu. Rev. Food Sci. Technol. 13(1):19–34. 10.1146/annurev-food-052720-112443.

Lim FS, González-Cabrera J, Keilwagen J et al. 2024. Advancing pathogen surveillance by nanopore sequencing and genotype characterization of Acheta domesticus densovirus in mass-reared house crickets. Sci Rep 14(1):8525. 10.1038/s41598-024-58768-3.

Llandres AL, Marques GM, Maino JL et al. 2015. A dynamic energy budget for the whole life-cycle of holometabolous insects. Ecol Monogr 85(3):353–371. 10.1890/14-0976.1.

Magara HJO, Niassy S, Ayieko MA et al. 2021. Edible crickets (Orthoptera) around the world: distribution, nutritional value, and other benefits—A review. Frontiers in Nutrition 7(10.3389/fnut.2020.537915.

Mahavidanage S, Fuciarelli TM, Li X et al. 2023. The effects of rearing density on growth, survival, and starvation resistance of the house cricket *Acheta domesticus*. J Orthoptera Res 32(1):25–31.

Mamai W, Lobb LN, Bimbilé Somda NS et al. 2018. Optimization of mass-rearing methods for *Anopheles arabiensis* larval stages: Effects of rearing water temperature and larval density on mosquito life-history traits. J Econ Entomol 111(5):2383–2390. 10.1093/jee/toy213.

Masaki S. 1978. ’Chapter 4 Seasonal and latitudinal adaptations in the life cycles of crickets’. In: Dingle H ed. *New York, NY*: *Springer US*. 72-100.

Mauritsson K, Jonsson T. 2023. A new flexible model for maintenance and feeding expenses that improves description of individual growth in insects. Sci Rep 13(1):16751. 10.1038/s41598-023-43743-1.

McFarlane JE. 1964. Factors affecting growth and wing polymorphism in *Gryllodes sigillatus* (Walk.): Dietary protein level and a possible effect of photoperiod. Can J Zool 42(5):767–771. 10.1139/z64-074.

Meister H, Esperk T, Välimäki P et al. 2017. Evaluating the role and measures of juvenile growth rate: latitudinal variation in insect life histories. Oikos 126(12):1726–1737. 10.1111/oik.04233.

Mirth CK, Anthony Frankino W, Shingleton AW. 2016. Allometry and size control: what can studies of body size regulation teach us about the evolution of morphological scaling relationships? Curr Opin Insect Sci 13:93–98. 10.1016/j.cois.2016.02.010.

Moore MP, Martin RA. 2019. On the evolution of carry-over effects. J Anim Ecol 88(12):1832–1844. 10.1111/1365-2656.13081.

Morales-Ramos JA, Rojas MG, Dossey AT et al. 2020. Self-selection of food ingredients and agricultural by-products by the house cricket, *Acheta domesticus* (Orthoptera:Gryllidae): A holistic approach to develop optimized diets. PLOS ONE 15(1):e0227400. 10.1371/journal.pone.0227400.

Morales-Ramos JA, Tomberlin JK, Miranda C et al. 2024. Rearing methods of four insect species intended as feed, food, and food ingredients: a review. J Econ Entomol:toae040. 10.1093/jee/toae040.

Morales-Ramos JA, Kay S, Rojas MG et al. 2015. Morphometric analysis of instar variation in *Tenebrio molitor* (Coleoptera: Tenebrionidae). Ann Entomol Soc Am 108(2):146–159. 10.1093/aesa/sau049.

Mutamiswa R, Tarusikirwa VL, Nyamukondiwa C et al. 2023. Thermal stress exposure of pupal oriental fruit fly has strong and trait-specific consequences in adult flies. Physiol Entomol 48(1):35–44. 10.1111/phen.12400.

Muzzatti MJ, McConnell E, Neave S et al. 2022. Fruitful female fecundity after feeding *Gryllodes sigillatus* (Orthoptera: Gryllidae) royal jelly. The Canadian Entomologist 154(1):e50. 10.4039/tce.2022.39.

Nakamura T, Ylla G, Extavour CG. 2022. Genomics and genome editing techniques of crickets, an emerging model insect for biology and food science. Curr Opin Insect Sci 50:100881. 10.1016/j.cois.2022.100881.

Nowosielski JW, Patton RL. 1965. Variation in the haemolymph protein, amino acid, and lipid levels in adult house crickets, *Acheta domesticus* L., of different ages. J Insect Physiol 11(3):263–270. 10.1016/0022-1910(65)90074-0.

Okada K, Pitchers WR, Sharma MD et al. 2011. Longevity, calling effort, and metabolic rate in two populations of cricket. Behav Ecol Sociobiol 65(9):1773–1778. 10.1007/s00265-011-1185-3.

Pinheiro JC, Bates D. 2009. Mixed-Effects Models in S and S-PLUS. Springer.

R Core Team. 2023. ’Chapter R: A language and environment for statistical computing’. 4.3. Vienna, Austria: R Foundation for Statistical Computing.

Raheem D, Carrascosa C, Oluwole OB et al. 2019. Traditional consumption of and rearing edible insects in Africa, Asia and Europe. Crit. Rev. Food Sci. Nutr. 59(14):2169–2188. 10.1080/10408398.2018.1440191.

Rapkin J, Jensen K, Archer CR et al. 2018. The geometry of nutrient space–based life-history trade-offs: Sex-specific effects of macronutrient intake on the trade-off between encapsulation ability and reproductive effort in decorated crickets. The American Naturalist 191(4):452–474. 10.1086/696147 10.1086/696147.

Ritchie MW, Provencher JF, Allison JE et al. 2024. The digestive system of a cricket pulverizes polyethylene microplastics down to the nanoplastic scale. Environ Pollut 343. 10.1016/j.envpol.2023.123168.

Robinson K, Duffield KR, Ramirez JL et al. 2024. MINIstock: Model for INsect Inclusion in sustainable agriculture: USDA-ARS’s research approach to advancing insect meal development and inclusion in animal diets. J Econ Entomol:toae130. 10.1093/jee/toae130.

Sakaluk SK, Oldzej J, Poppe CJ et al. 2019. Effects of inbreeding on life-history traits and sexual competency in decorated crickets. Anim Behav 155:241–248. 10.1016/j.anbehav.2019.05.027.

Schindelin J, Arganda-Carreras I, Frise E et al. 2012. Fiji: an open-source platform for biological-image analysis. Nat. Methods 9(7):676–682. 10.1038/nmeth.2019.

Siddiqui SA, Aidoo OF, Ghisletta M et al. 2024. African edible insects as human food -a comprehensive review. Journal of Insects as Food and Feed 10(1):51–78. 10.1163/23524588-20230025.

Sorjonen JM, Valtonen A, Hirvisalo E et al. 2019. The plant-based by-product diets for the mass-rearing of *Acheta domesticus* and *Gryllus bimaculatus*. PLOS ONE 14(6):e0218830. 10.1371/journal.pone.0218830.

Stahlschmidt ZR, Choi J, Choy B et al. 2023. A simulated heat wave—but not herbicide exposure—alters resource investment strategy in an insect. J Therm Biol 116(103670. 10.1016/j.jtherbio.2023.103670.

Tedeschi LO, Greenwood PL, Halachmi I. 2021. Advancements in sensor technology and decision support intelligent tools to assist smart livestock farming. J Anim Sci 99(2):skab038. 10.1093/jas/skab038.

Tennessen Jason M, Thummel Carl S. 2011. Coordinating growth and maturation — Insights from *Drosophila*. Curr Biol 21(18):R750–R757. 10.1016/j.cub.2011.06.033.

Tomberlin JK, Picard CJ, Jordan HR et al. 2022. Government and industry investment plays crucial role in further establishment, evolution, and diversification of insect agriculture: a case example from the United States. Journal of Insects as Food and Feed 8(2):109–111. 10.3920/JIFF2022.x001.

Toms RB. 1993. More winged females of the cricket *Gryllodes supplicans* (Walker). S Afr J Zool 28(2):122–124. 10.1080/02541858.1993.11448304.

UN. 2015. Transforming our world: the 2030 Agenda for Sustainable Development.

Urquhart-Cronish M, Sokolowski MB. 2014. Gene-environment interplay in *Drosophila melanogaster*: Chronic nutritional deprivation in larval life affects adult fecal output. J Insect Physiol 69:95–100. 10.1016/j.jinsphys.2014.06.001.

van Huis A. 2020. Insects as food and feed, a new emerging agricultural sector: a review. Journal of Insects as Food and Feed 6(1):27–44. 10.3920/JIFF2019.0017.

van Huis A, Rumpold B, Maya C et al. 2021. Nutritional qualities and enhancement of edible insects’. In: Stover PJ, Balling R eds. 551-576.

Van Peer M, Berrens S, Coudron C et al. 2024. Towards good practices for research on *Acheta domesticus*, the house cricket. Journal of Insects as Food and Feed:1–17. 10.1163/23524588-00001042.

Vogel M, Shah PN, Voulgari-Kokota A et al. 2022. Health of the black soldier fly and house fly under mass-rearing conditions: innate immunity and the role of the microbiome. Journal of Insects as Food and Feed 8(8):857–878. 10.3920/jiff2021.0151.

Wenning MJ, Piotrowski T, Janzen J et al. 2022. Towards monitoring of a cricket production using instance segmentation. Journal of Insects as Food and Feed 8(7):763–772. 10.3920/JIFF2021.0165.

Woodman JD. 2012. Cold tolerance of the Australian spur-throated locust, *Austracris guttulosa*. J Insect Physiol 58(3):384–390. 10.1016/j.jinsphys.2011.12.015.

Zhang W, Chang X-Q, Hoffmann A et al. 2015. Impact of hot events at different developmental stages of a moth: the closer to adult stage, the less reproductive output. Sci Rep 5(1):10436. 10.1038/srep10436.

