## Supplementary Materials for "Growth, development, and life history of a mass-reared edible insect, *Gryllodes sigillatus* (Orthoptera: Gryllidae)"

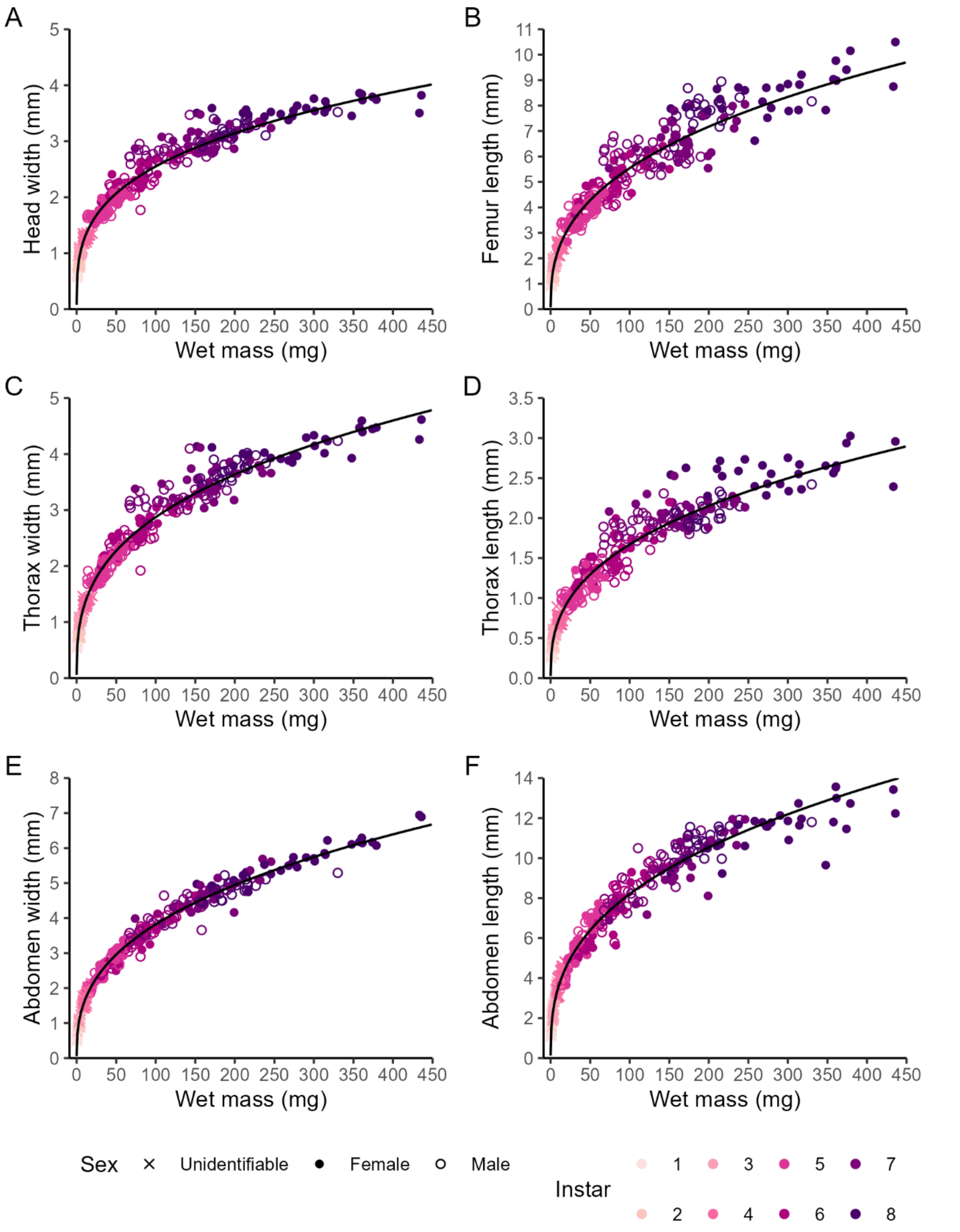


**Fig. S1**. A) Head width, B) Femur length, C) Thorax width, D) Thorax length, E) Abdomen width, and F) Abdomen length of 355 *Gryllodes sigillatus* crickets by wet mass. Colors denote instar. Crosses denote juvenile crickets of unidentifiable sex (instars 1 – 4), closed circles denote female crickets (instars 5 – 8), and open circles denote male crickets (instars 5 – 8). Solid black line indicates the fitted linear regression to $Log_{10}$ transformed traits and $Log_{10}$ transformed wet mass, pooling sex (Table S1).

**Table S1**. Regression equations and Coefficient of Determination, $R^2$ (%), for the allometric relationships shown in **Fig.** S1.

| **Figure panel** | **Trait** | **Equation** | **$R^2$ (%)** |
|---|---|---|---|
| Fig. S1A | Head width (mm) by wet mass (mg) | $Log_{10}$(Head width) = 0.30 × $Log_{10}$(Wet mass)– 0.20 | 97.26 |
| Fig. S1B | Femur length (mm) by wet mass (mg) | $Log_{10}$(Femur length) = 0.37 × $Log_{10}$(Wet mass)– 00009 | 96.67 |
| Fig. S1C | Thorax width (mm) by wet mass (mg) | $Log_{10}$(Thorax width) = 0.34 × $Log_{10}$(Wet mass)– 0.22 | 97.79 |
| Fig. S1D | Thorax length (mm) by wet mass (mg) | $Log_{10}$(Thorax length) = 0.36 × $Log_{10}$(Wet mass)– 0.51 | 96.54 |
| Fig. S1E | Abdomen width (mm) by wet mass (mg) | $Log_{10}$(Abdomen width) = 0.37 × $Log_{10}$(Wet mass) – 0.16 | 98.92 |
| Fig. S1F | Abdomen length (mm) by wet mass (mg) | $Log_{10}$(Abdomen length) = 0.36 × $Log_{10}$(Wet mass) + 0.19 | 97.78 |

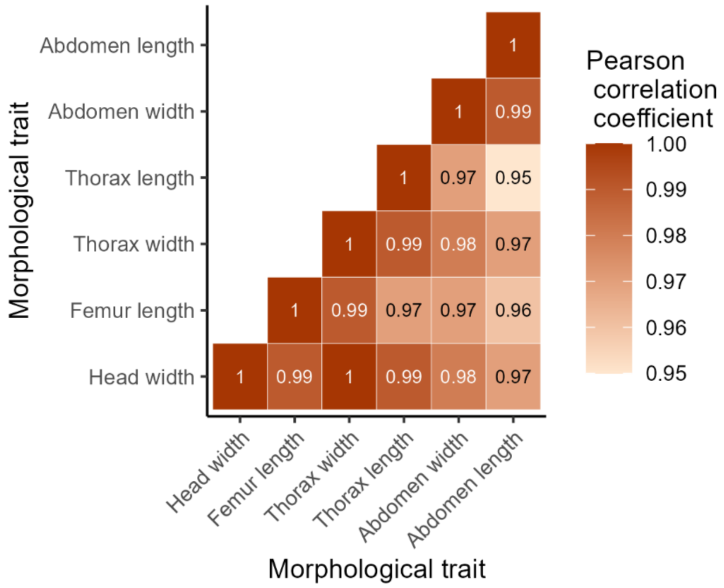


**Fig. S2**. Pearson correlation coefficients (colors) for pairwise correlations between morphological traits throughout ontogeny, pooling sex (n = 355 crickets).

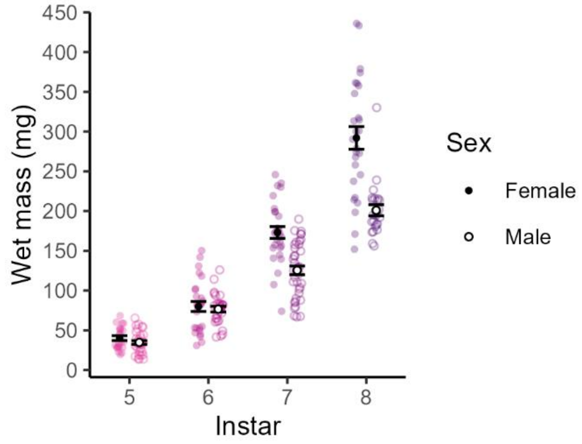


**Fig. S3**. Wet mass of male (open circles) and female (closed circles) *Gryllodes sigillatus* crickets (n = 231) for instars 5-8 (colors) when sex could be determined.
